# Canonical and noncanonical contribution of thyroid hormone receptor isoforms alpha and beta to cardiac hypertrophy and heart rate in male mice

**DOI:** 10.1101/2023.11.24.568041

**Authors:** Daniela Geist, G. Sebastian Hönes, Susanne C. Grund, Janina Pape, Devon Siemes, Philippa Spangenberg, Elen Tolstik, Stefanie Dörr, Nadine Spielmann, Helmut Fuchs, Valerie Gailus-Durner, Martin Hrabe de Angelis, Jens Mittag, Daniel R. Engel, Dagmar Führer, Kristina Lorenz, Lars C. Moeller

## Abstract

**Background:** Stimulation of ventricular hypertrophy and heart rate are two major cardiac effects of thyroid hormone (TH). Aim of this study was to determine *in vivo* which TH receptor (TR), α or β, and which mode of TR action, canonical gene expression or DNA-binding independent noncanonical action, mediate these effects.

**Material and methods:** We compared global TRα and TRβ knockout mice (TRα^KO^; TRβ^KO^) with WT mice to determine the TR isoform responsible for T3 effects. The relevance of TR DNA-binding was studied in mice with a mutation in the DNA-binding domain that selectively abrogates DNA binding and canonical TR action (TRα^GS^; TRβ^GS^). Hearts were studied with echocardiography at baseline and after seven weeks T3-treatment. Gene expression was measured with real-time PCR. Heart rate was recorded with radiotelemetry transmitters for seven weeks in untreated, hypothyroid and T3-treated mice.

**Results:** T3 induced ventricular hypertrophy in WT and TRβ^KO^ mice, but not in TRα^KO^ mice. Hypertrophy was also induced in TRα^GS^ mice. Thus, hypertrophy is mostly mediated by noncanonical TRα action. Similarly, repression of *Mhy7* occurred in WT and TRα^GS^ mice. Basal heart rate was largely dependent on canonical TRα action. But responsiveness to hypothyroidism and T3-treatment as well as expression of pacemaker gene *Hcn2* were still preserved in TRα^KO^ mice, demonstrating that TRβ could compensate for absence of TRα.

**Conclusion:** T3-induced cardiac hypertrophy could be attributed to noncanonical TRα action, whereas heart rate regulation was mediated by canonical TRα action. TRβ could substitute for canonical, but not noncanonical TRα action.

## Introduction

The heart is a major target organ of thyroid hormones (TH) and especially heart rate and ventricular growth are stimulated by TH. TH effects are mediated by the TH receptors (TR) TRα1 and TRβ1 and 2 (1-3). TRs act as ligand-dependent transcription factors regulating the expression of TH-target genes by binding to thyroid hormone response elements (TRE) on the DNA (type 1 TR signaling, canonical). In addition, T3 (3,3’,5-triiodo-l-thyronine) and TRs mediate activation of signaling pathways, e.g. PI3K/AKT and MAPK/ERK (type 3 TR signaling, noncanonical) (4-11). This mode of action is independent from DNA binding of TRs. The precise mechanisms of TH action leading to different TH-mediated cardiac effects is still unclear and requires the determination of the responsible TR isoform, TRα or TRβ, and of the mode of TR action, canonical or noncanonical. TH excess, either hyperthyroidism in patients or TH treatment of mice, leads to cardiac hypertrophy through a combination of TH action in cardiomyocytes, including regulation of gene expression (e.g. *Myh6* and *7*, adrenergic receptors) and regulatory pathway activation (e.g. PI3K and MPAPK signaling), and interaction with other signaling systems, especially the sympathetic nervous system (12). Although TRα is the predominant TR isoform in the heart, T3-induced ventricular growth has not been unanimously attributed to a TR isoform. Thyroxine (T4) treatment of hypothyroid mice for four weeks led to an increase in cardiac mass in wild-type (WT) and TRα knockout mice, whereas absence of TRβ in TRβ knockout mice prevented the T4-induced increase in cardiac mass (13). Furthermore, T3 induced cardiac hypertrophy only in WT, but not in mice expressing a dominant-negative TRβ^Δ337T^ mutant receptor (14, 15). Thus, TH-induced cardiac hypertrophy was attributed to TRβ.

Interestingly, T4 treatment in rats increased heart weight and Akt phosphorylation, indicating activation of the PI3K by TH (16). In mice, T4 treatment induced cardiac hypertrophy, which was abrogated by co-treatment with rapamycin, an mTOR inhibitor (17). *In vitro*, T3 rapidly increased PI3K activity and subsequently Akt and mTOR phosphorylation in rat cardiomyocytes (18). In addition, TRα and the regulatory PI3K subunit p85α formed complexes in cardiomyocytes. Together, these data suggested that ventricular growth in response to TH could at least in part be mediated by noncanonical TR signaling.

Here we studied genetic mouse models to determine which TR, TRα or TRβ, and which mechanism, canonical or noncanonical signaling, mediates TH-induced cardiac effects such as ventricular growth or heart rate regulation. We compared global TRα knockout mice (TRα^KO^) and TRβ knockout mice (TRβ^KO^) with WT mice to determine the TR isoform responsible for the T3 effects on the on cardiac hypertrophy and heart rate. To determine the relevance of TRs’ DNA-binding and, thus, the potential relevance of noncanonical TR signaling, we studied mice with mutations in the TR DNA-binding P-box, changing glutamic acid (E) and glycine (G) to glycine and serine (S) (EG to GS, codons 71/72 in *Thra* and 125/126 in *Thrb*, respectively) (7). These mutations abrogate the TRs’ DNA-binding and canonical actions in TRα^GS^ and TRβ^GS^ mice. A comparison of WT, TRα^KO^ and TRβ^KO^ and TRα^GS^ or TRβ^GS^ mice allows to attribute TH effects to the relevant TR isoform and to the underlying TR signaling mechanism. The results demonstrated that T3-induced ventricular growth was mediated by noncanonical TRα action and heart rate was regulated by canonical TRα action with contribution of TRβ.

## Materials and Methods

### Animal studies

All animal experiments were approved by the Landesamt für Natur, Umwelt und Verbraucherschutz Nordrhein-Westfalen (LANUV-NRW) and performed in accordance with the German regulations for Laboratory Animal Science (GV SOLAS) and the European Health Law of the Federation of Laboratory Animal Science Associations (FELASA). TRα knockout (TRα^0/0^) and TRβ knockout (TRβ^-/-^) mice, here referred to as TRα^KO^ and TRβ^KO^, respectively, were obtained from the European Mouse Mutant Archive (https://www.infrafrontier.eu) (2, 3). Generation of TRα^GS^ and TRβ^GS^ mutant mice was described previously (7). Mice were housed in the central animal facility at the University Hospital Essen in individually ventilated cages at 21±1 °C in an alternating 12:12 hour light-dark cycle and fed standard chow (Sniff, Soest, Germany) and tap water provided *ad libitum*. For echocardiography studies T3 was administered orally via drinking water containing 400 ng/mL T3 (Sigma-Aldrich, USA) for seven weeks. Echocardiography measurements were performed with mice aged approximately eight weeks (echo 1) and 7 weeks later (echo 2) (Figure 1A). Mice were anesthetized with pentobarbital (20 mg/g body weight) and examined with a Vevo2100 (VisualSonics) echocardiography device. Fractional shortening, end-diastolic interventricular septum thickness (IVS;d), left ventricular posterior wall thickness (LVPW;d) and left ventricular inner dimension (LVID;d) were recorded and calculated (19, 20).

**Figure 1:**
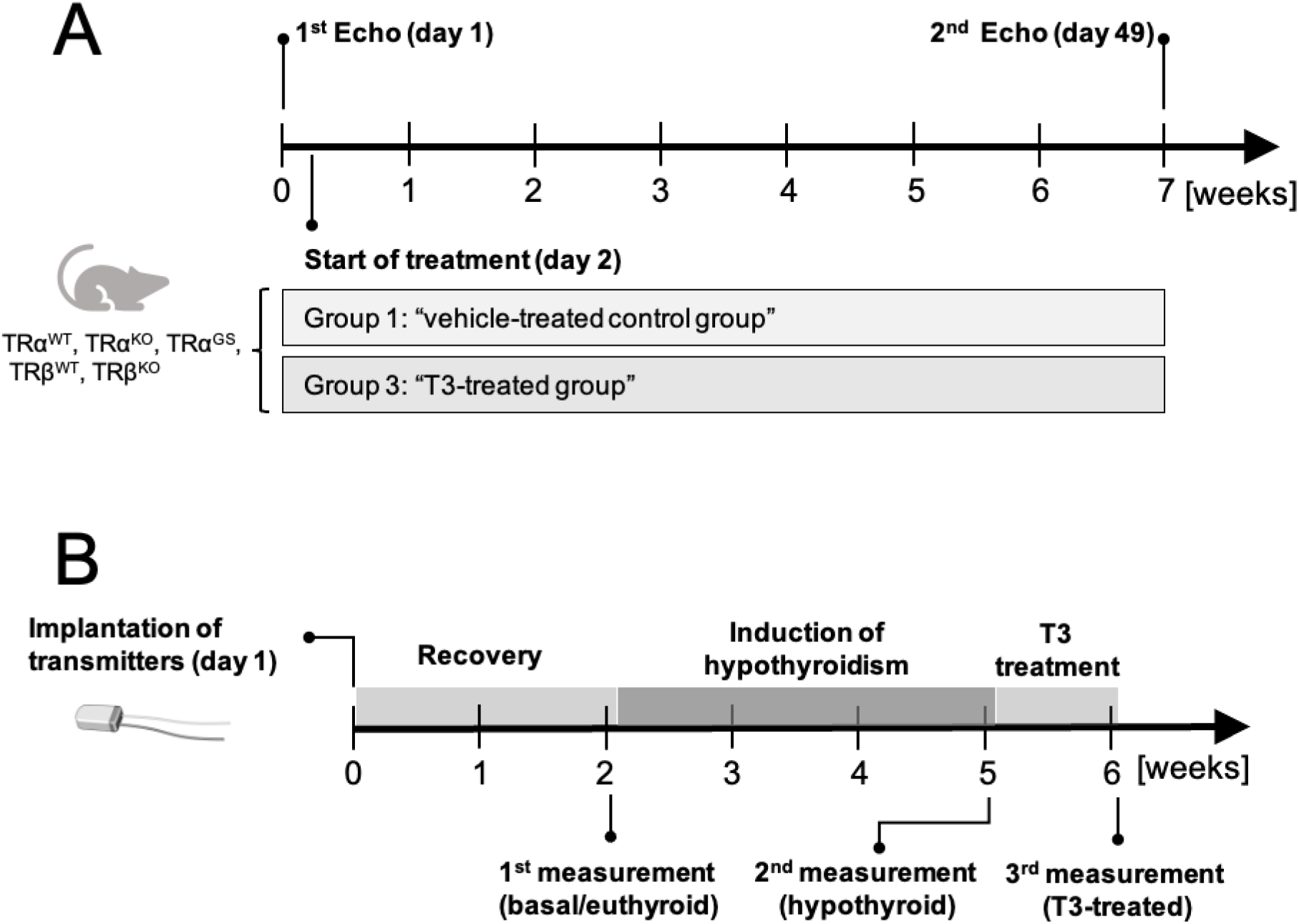
Echocardiography and radio telemetry study design. (A) Two months old male mice were studied with echocardiography on day 1 (Echo 1). From day 2 on, mice received T3 (400 ng/mL) or vehicle (9.3 µM NaOH; control group) in the drinking water. Echo 2 was performed after 7 weeks. At the end of the experiment, hearts and tibias were collected. (B) Experimental setup for radio telemetry experiments. This study was performed over a time span of 6 weeks. After surgery, mice were treated with analgesic for four days during the recovery phase and allowed to further recover for a total of 10-14 days. Afterwards, mice were measured in all three thyroid hormone states: euthyroid (3 days), hypothyroid (3 days) and T3-treated (6 days, divided into early and late phase).

As part of the systemic phenotyping, electrocardiography (ECG) was performed at the German Mouse Clinic (GMC) in Munich in awake 15-weeks-old untreated WT, TRα^KO^, TRα^GS^, TRβ^KO^, TRβ^GS^ mice (21, 22) Mice were maintained in IVC cages with water and standard mouse chow according to the directive 2010/63/EU, German laws and GMC housing conditions (www.mouseclinic.de). All tests were approved by the responsible authority of the district government of Upper Bavaria.

For radio telemetry experiments, chronic hypothyroidism was induced by feeding mice low iodine food (LID; TD.95007, Harlan Laboratories) and supplementing the drinking water with 0.04% methimazole, 0.5% sodium perchlorate (ClO4-) and 0.3% saccharine as sweetener for 3 weeks.

High T3 serum concentrations were achieved by adding 400 ng/mL T3 to drinking water (Sigma-Aldrich, USA) for 6 days.

Long-term *in vivo* heart rate was measured with a radio telemetry system (Data Science International) (23). Transmitters (DSI PhysioTel® Transmitter (ETA-F10)) were implanted in 4– 5-month-old mice with a minimal body weight of 25 g. Mice were anesthetized with fully antagonizable anesthesia followed by a subcutaneous injection of carprofen (5 mg/kg BW). After placing the transmitter into the abdominal cavity, the anode was tunneled subcutaneously to the neck, where the loop was attached to a muscle with surgical thread. The cathode was attached to a muscle below the heart so that both electrodes formed a diagonal line over the heart. Following surgery, mice were woken up with the anesthesia antagonist and supplied with 15% glucose solution as drinking water and steeped food pellets. Mice were allowed to recover for 10-14 days and were treated with analgesics for four days. After the recovery period, heart rate was constantly recorded for three days each in untreated mice and after treating mice with hypothyroidism-inducing food and drinking water for 3 weeks (Fig. 1B). Next, T3 was added to the drinking water and heart rate was recorded for 6 days to investigate the transition from hypothyroidism to early (day 1-3) as well as late T3 treatment phase (day 4-6). For analysis of the recorded data, they were assigned to the natural active and inactive state of the animals. Mean values were saved every 30 seconds.

### Organ collection

Mice were sacrificed by cervical dislocation and body weight was recorded. Blood was collected by heart puncture and mice were perfused with PBS containing 5 U/mL heparin, followed by a second perfusion with PBS. Hearts were obtained, atria were removed, and ventricles were weighed. The upper half of hearts was stored in paraformaldehyde and the apex was cut in small pieces, frozen in liquid nitrogen and stored at -80 °C. Tibia length was measured for normalization of heart weights (24). Blood was collected in Microvette® tubes (Sarstedt, Germany), stored on ice for 30 min to induce coagulation and centrifuged at 4 °C with 17,000×g for 25 min. Serum was used for the determination of TH levels.

### Gene expression analysis

Total RNA was isolated from 20-30 mg heart tissue (QIAshredder and RNeasy Mini Kit, Qiagen) and eluded in 20 µL RNase-free water. Total RNA concentration was measured with a NanoDrop2000. RNA integrity was tested on a 1.2% agarose gel (RNA-free water). Samples were incubated at 65 °C for 10 min before being loaded on the agarose gel and visualized at a Molecular Imager® VersaDoc™. 1 µg of total RNA was transcribed into cDNA by using a SuperScript™ III Reverse Transcriptase kit (Invitrogen) and random hexamer primers. Gene expression was normalized to that of three reference genes (*18S*, *Gapdh* and *Pol2a2*). For analysis and calculation of gene expression only Ct values below 35 cycles were evaluated using the efficiency corrected method (25). Primer sequences are listed in Table S1.

### Thyroid hormone serum concentration

TH serum concentrations were measured with commercial FT3, FT4, and TT4 ELISA kits (DRG Instruments GmbH, Marburg) on a VersaMax Microplate Reader (Molecular Devices, Biberach). Minimum detectable levels of TH were 0.5 μg/dL for TT4, 0.05 ng/dL for FT4 and 0.05 pg/mL for FT3.

### Statistics and software

Data were analyzed with GraphPad Prism 6 (GraphPad, San Diego, USA) and presented mean ± standard error of mean (SEM). To test for significance, we used one-way ANOVA with Tukey‘s multiple comparison test for comparison of all groups, Sidak’s multiple comparison test for comparison of selected groups and Dunnett’s multiple comparison test for comparing all groups to one reference group. Two-way ANOVA with Bonferroni‘s *post hoc* correction was used for echocardiography evaluation. Multiple t-test was used to compare hourly averaged heart rate values measured over three days. Differences were considered significant with p<0.05.

## Results

### T3 treatment induces ventricular growth via noncanonical TRα signaling

We compared posterior and interventricular wall diameters (LVPW and IVS) of male TRα^KO^, TRα^GS^ and TRβ^KO^ mice with their respective WT littermates with echocardiography before and after 7 weeks of T3 treatment (400 ng/mL), starting at an age of 8 to 10 weeks. T3 treatment for seven weeks led to 1.5 to 2.0-fold increased FT3 serum concentration in all genotypes (Figure S1A). FT4 concentrations were below detection limit and TT4 was significantly reduced as well due to T3 treatment (Figure S1B).

Already at baseline, TRα^KO^ mice had smaller and lower absolute and relative heart weights compared to their WT littermates, whereas hearts of TRβ^KO^ mice did not differ from WT mice (Figure 2A-D). In WT mice, seven weeks of T3 treatment led to an increase in LVPW and IVS (WT littermates of TRα and TRβ mutant strains) (Figure 2A,B). In contrast, T3 treatment did not induce an increase in wall diameters in TRα^KO^ mice and their LVPW and IVS dimensions remained smaller than those of WT mice, indicating that absence of TRα prevented cardiac hypertrophy. A similar increase in LVPW as in WT mice was found in TRβ^KO^ mice and the IVS and LVPW diameters were not different from their WT littermates, demonstrating that absence of TRβ had no major influence on T3-induced cardiac hypertrophy. Unlike in TRα^KO^ mice the cardiac wall diameters of TRα^GS^ mice were increased by T3 treatment, although the hypertrophic response was less pronounced than in WT controls. These data indicate that T3/TRα-mediated cardiac hypertrophy did not depend on canonical TRα signaling and was rather mediated by noncanonical TRα action.

**Figure 2:**
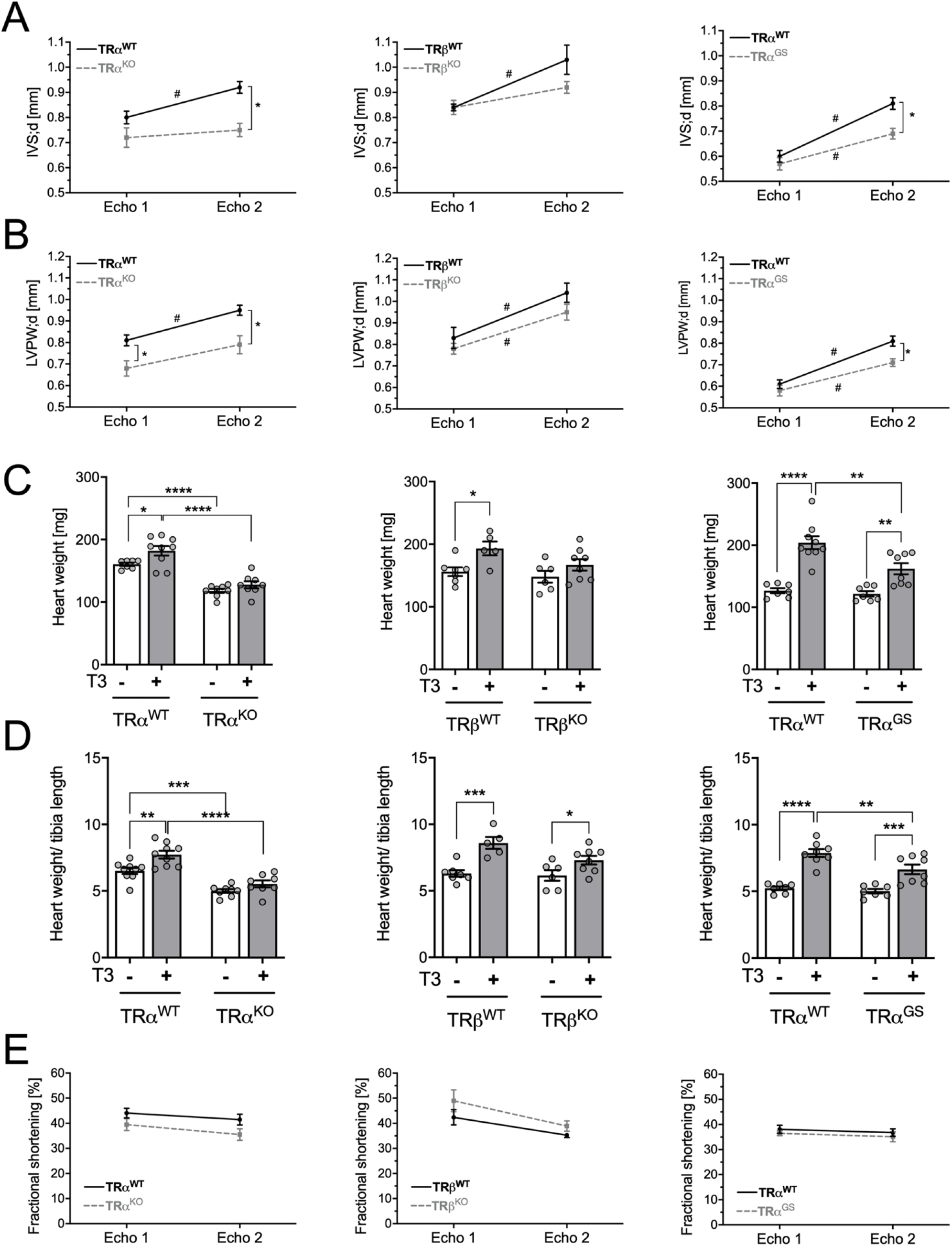
Echocardiography parameters: baseline and after T3-treatment of mice. Echocardiography measurements of male TRα^WT^, TRα^KO^, TRα^GS^, TRβ^WT^ and TRβ^KO^ mice were conducted at experimental day 1 and after 7 weeks of T3 treatment. (A) Interventricular septum thickness (IVS;d) and (B) left ventricular posterior wall thickness (LVPW;d) were determined to analyze heart wall thickness in TR mutant mice and their WT littermates (TRα^KO^ n=7-9, TRα^GS^ n=7-9, TRβ^KO^ n=5-7; TRα^KO^ Two-way repeated ANOVA with Bonferroni‘s multiple comparison test. Values are mean ± SEM); *P<0.05; intra-group comparison between Echo1 and Echo2; #P<0.05) (C) *Ex vivo* heart weight (ventricles) was determined after the last echocardiography. (D) Heart weight was normalized to tibia length to calculate the extent cardiac hypertrophy (TRα^KO^ n=7-9, TRα^GS^ n=7-9, TRβ^KO^ n=5-7; mean ± SEM; One-way ANOVA with Tukey‘s multiple comparison test; *P<0.05, **P<0.01, ***P<0.001, ****P<0.0001). (E) Heart weight was normalized to tibia length to calculate the extent cardiac hypertrophy. (TRα^KO^ strain n=7-9, TRα^GS^ strain=7-9, TRβ^KO^ strain n=5-7; mean ± SEM; Two-way repeated ANOVA with Bonferroni‘s multiple comparison test). Values are mean ± SEM.

At the end of the experiment, ventricular weight and tibia length were measured. The ventricular weight (Figure 2C) as well as the ventricular weight/tibia length ratio (Figure 2D) was increased by T3 treatment in all WT littermates compared to untreated WT mice. Ventricular weight/tibia length ratio was also increased in TRβ^KO^ mice but not in TRα^KO^ mice (Figure 2C,D), confirming that ventricular growth by T3 is mediated by TRα. Interestingly, the ventricular weight to tibia length ratio was also increased in TRα^GS^ mice (Figure 2C,D), further demonstrating that DNA-binding of TRα was not required for this effect.

To assess cardiac function in response to seven weeks of T3 treatment, fractional shortening was measured as a parameter for systolic function. Fractional shortening was not influenced by T3 in this experimental setting regardless of the genotype (Figure 2E).

In accordance with increased echocardiographic wall diameters and ventricular weight/tibia length ratio, T3 treatment led to cardiomyocyte hypertrophy in WT and TRβ^KO^ mice, which was absent in TRα^KO^ mice (Figure 3). Again, the hypertrophic T3-effect was preserved in TRα^GS^ mice, demonstrating that T3-induced cardiac growth is a consequence of cardiomyocyte hypertrophy and independent from TRα DNA-binding.

**Figure 3:**
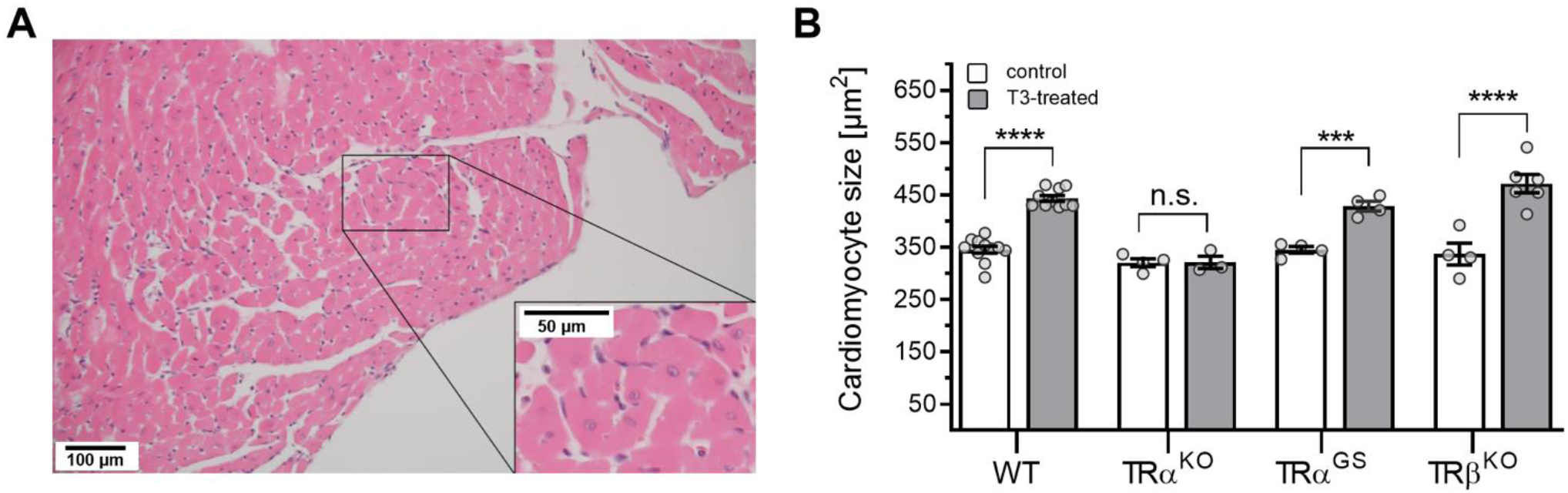
Cardiomyocyte size in untreated and T3-treated WT, TRα^KO^, TRα^GS^ and TRβ^KO^ hearts. (A) Heart sections of control and T3-treated mice were stained with hematoxylin and eosin and examined via light microscopy. Image shows H&E-stained heart section from a control WT mouse. (B) Cardiomyocyte size in male control (white) and T3-treated (grey) mice was measured and calculated with ImageJ. A minimum of 500 cells was evaluated per group. Scale bars represent 100 µm and 50 µm, respectively. (n=3-6; mean ± SEM; One-way ANOVA with Sidak‘s multiple comparison test; control vs. T3-treated within one genotype; ***P<0.01, ****P<0.0001; n.s.= not significant.)

### T3-mediated cardiac gene induction is regulated by TRα and TRβ

Next, we measured gene expression in control and T3-treated hearts from all genotypes. *Thra* and *Thrb* expression was absent or downregulated in respective TR^KO^ hearts (Figure 4). T3 treatment resulted in downregulation of *Thra* in TR WT and similarly in TRβ^KO^ and TRα^GS^ hearts (Figure 4). *Myh6* expression was not different between the genotypes and largely unaltered by T3 with only a mild increase in TRα^KO^ mice (Figure 4). Compared to WT, basal *Myh7* expression was increased in TRα^KO^ and TRα^GS^ mice, compatible to our previous observations (7) (Figure 4). *Mhy7* was repressed by T3 treatment in WT mice. In TRβ^KO^ mice, basal *Myh7* expression and T3-mediated repression were not different from WT, demonstrating the TRα WT effect. Strikingly, complete *Myh7* repression by T3 also occurred in TRα^KO^ and TRα^GS^ mice (Figure 4), which strongly suggests that T3-mediated *Myh7* repression does not solely depend on TRα and can also be mediated by TRβ.

**Figure 4:**
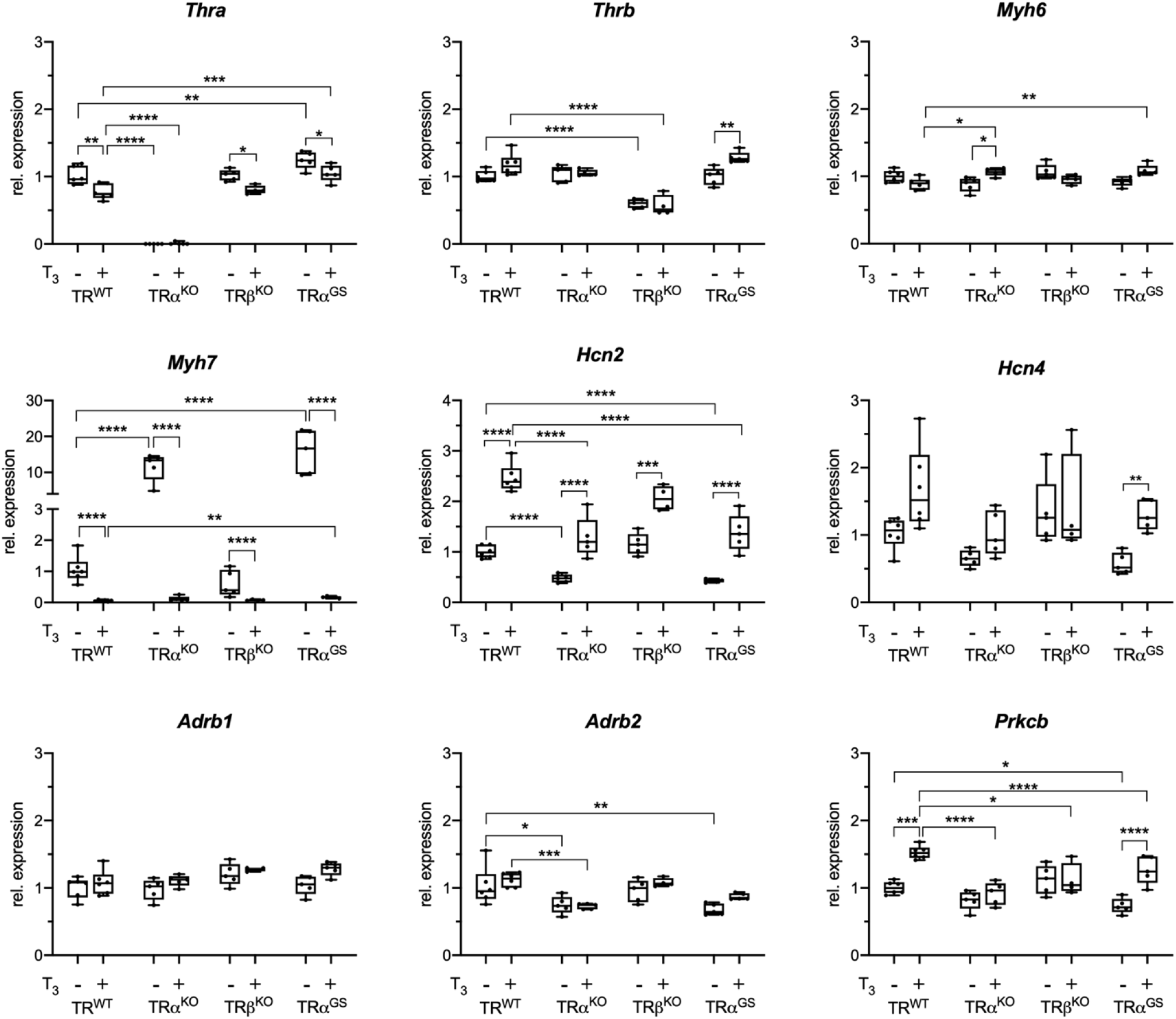
Relative gene expression in hearts of control and T3-treated mice. Relative gene expression of TH receptor α (*Thra*), TH receptor β (*Thrb*), myosin heavy chain 6 (*Myh6*) and 7 (*Myh7*), activated cyclic nucleotide gated potassium channel 2 (*Hcn2*) and 4 (*Hcn4*), adrenergic receptor 1 (*Adrb1*) and 2 (*Adrb2*) (n=5-6; one-way ANOVA with Sidak‘s multiple comparison test (Comparisons are male control vs. T3-treated and TR^WT^ vs. TRα^KO^, TRβ^KO^ and TRα^GS^, respectively; values are shown as boxplots with whiskers (min to max); *P<0.05, **P<0.01, ***P<0.001, ****P<0.0001.)

Next, we measured the expression of T3-responsive pacemaker ion channels *Hcn2* and *Hcn4* that contribute to regulation of heart rate. Basal *Hcn2* expression was reduced in TRα^KO^ and TRα^GS^ mice. Interestingly, *Hcn2* induction by T3 treatment was preserved in TRα^KO^ and TRα^GS^ mice, which, similar to *Myh7* repression, demonstrates that TRα action is not required for *Hcn2* induction. Basal *Hcn2* expression and T3-mediated induction in TRβ^KO^ mice were the same as in WT mice, demonstrating the effect of the WT TRα. *Hcn4* expression in the genotypes and basal or with T3 treatment showed a similar pattern, although not significant except for TRα^GS^, which still showed that DNA-binding of TRα is not required for *Hcn4* induction. Expression of *Adrb1* and *2* was not responsive to T3 (Figure 4). Protein Kinase C Beta (*Prkcb*) expression was induced by T3 in WT, but not in TRα^KO^ mice. While dependent on TRα, it seemed not to be the canonical mechanism that leads to *Prkcb* induction as it was also induced by T3 in hearts of TRα^GS^ mice.

### Heart rate and its changes by TH serum concentrations are controlled by canonical TRα signaling

ECGs were recorded in conscious animals and cardiac electrical activity was detected non-invasively through the animals’ paws. Heart rate was reduced in TRα^KO^ mice compared to WT controls (Figure 5A). Heart rates of TRβ^KO^ and TRβ^GS^ mice, genotypes with intact TRα signaling, were not different from those of their respective WT littermates (Figure 5A). Heart rate is therefore regulated by TRα, a classical, long known physiological effect (26).

**Figure 5:**
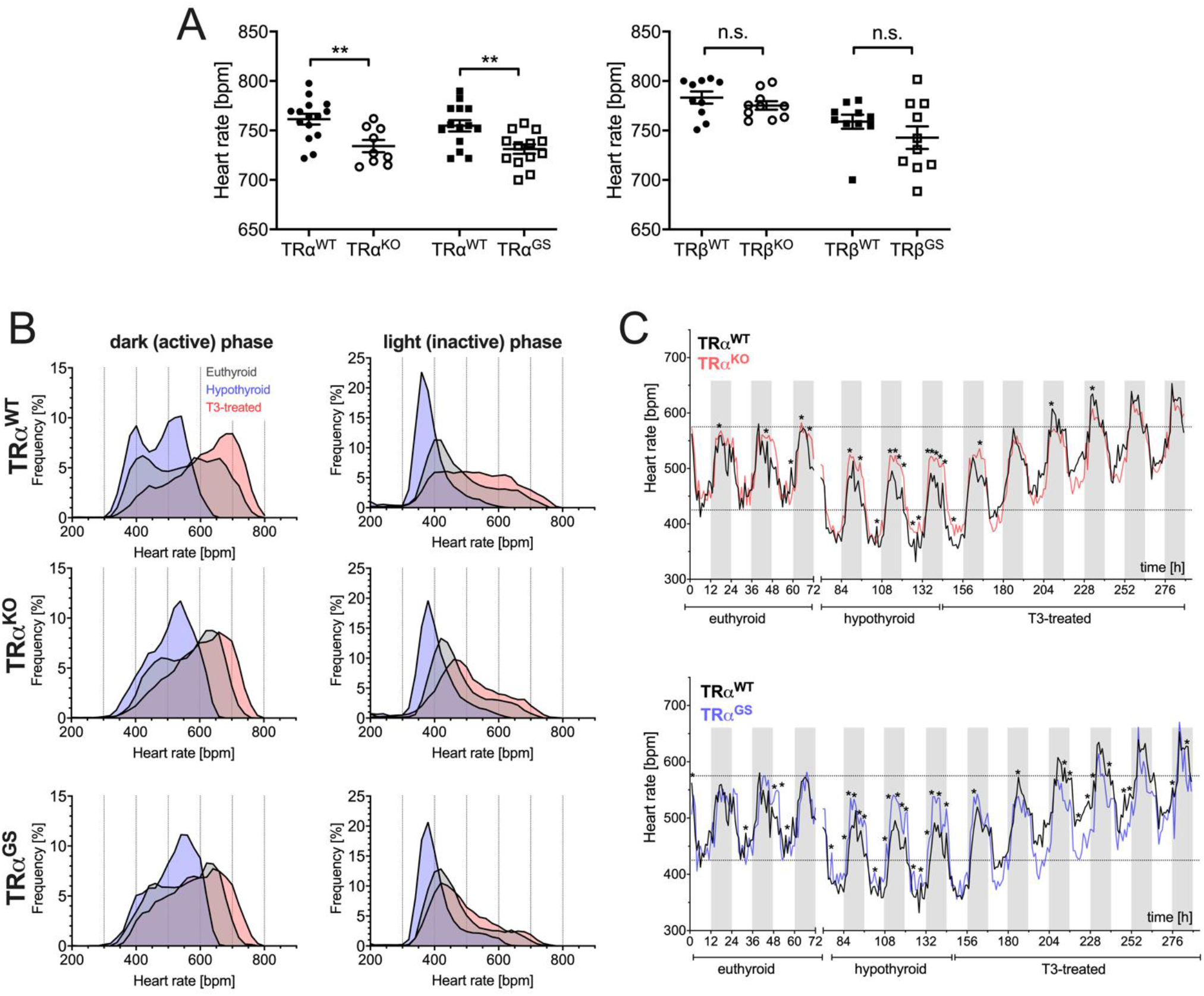
Heart rate is regulated by canonical TRα action with contribution from TRβ. (A) Heart rate was analyzed via electrocardiography in conscious euthyroid male WT, TRα^KO^ and TRα^GS^ (left panel, n=9-15) and TRα^WT^, TRβ^KO^ and TRβ^GS^ (right panel, n=10) mice at the age of 15 weeks. Mean ± SEM; One-way ANOVA with Sidak‘s multiple comparison test, TRα^WT^ vs. TRα^KO^ and TRα^GS^ respectively and TRβ^WT^ vs. TRβ^KO^ and TRβ^GS^ respectively; **P<0.01; n.s.= not significant. (B) Heart rate was continuously recorded via radio telemetry for three days in freely-moving euthyroid (grey), hypothyroid (blue) and early and late T3-treated (red) male TRα^WT^, TRα^KO^ and TRα^GS^ mice. (n=6-8). Relative frequency of heart rates values was calculated in 20 bpm steps and expressed as %. (C) Effect of TH status on heart rate male WT, TRα^KO^ and TRα^GS^ mice (n=6-8). Upper penal: Comparison of TRα^WT^ (black curve) and TRα^KO^ (red curve) mice. Lower panel: Comparison of TRα^WT^ (black curve) and TRα^GS^ (blue curve) mice. Multiple t- test calculated for hourly means of TRα^WT^ vs. TRα^KO^ or TRα^GS^, respectively; *P<0.05. Heart rate was recorded for three days in eu-, hypothyroid and early and late T3 treatment phase of male TRα^WT^, TRα^KO^ and TRα^GS^ mice. Upper panel: Comparison of TRα^WT^ (black) and TRα^KO^ (red). Lower panel:) and TRα^GS^ (blue) mice.

To analyze the response of heart rate in the TRα mutant mice to hypothyroidism and T3 treatment, we implanted telemetry devices into 4–5-month-old male mice for long-time monitoring of heart rates in conscious animals (WT, TRα^KO^ and TRα^GS^ mice).

In WT mice, hypothyroidism resulted in lower heart rates and T3 treatment in higher heart rates (Figure 5B). These changes in heart rate distribution, due to low or high TH, were more apparent in the active phase. Interestingly, the decrease and increase of heart rate in response to hypothyroidism or T3 treatment, respectively, was much less pronounced in TRα^KO^ and TRα^GS^ mice, especially in the active phase (Figure 5B). During the inactive phase there was only a mild adaptation due to hypothyroidism towards lower heart rates. Radiotelemetry recordings of TRα^WT^ vs. TRα^KO^ and TRα^WT^ vs. TRα^GS^ during the course of the protocol (untreated, methimazole/low iodine fed and T3-treated) show that the deflection of heart rate was significantly reduced in TRα^KO^ and TRα^GS^ mice (Figure 5C). These data demonstrate that the response of heart rate to the TH concentration, hypothyroidism or T3 treatment, is mainly regulated by canonical TRα signaling. Interestingly, heart rate was still modulated in TRα^KO^ mice (Figure 5B,C), demonstrating that TRβ contributes to heart rate regulation by TH.

## Discussion

Main results from phenotyping untreated and T3-treated WT, TRα^KO^, TRα^GS^ and TRβ^KO^ mice were that cardiac hypertrophy could be attributed to noncanonical TRα action, whereas heart rate regulation was mediated by canonical TRα action. As TH still modulated gene expression and heart rate in TRα^KO^ mice, TRβ contributed to or even partially substituted for canonical TRα action.

### Cardiac hypertrophy is a noncanonical TRα effect

TH induced cardiac hypertrophy in patients with untreated hyperthyroidism and in rodent models (13-15, 17, 27-29). While some mouse studies attributed this effect to TRβ action, there was also evidence of PI3K signaling pathway activation by TH, although this effect was not directly attributed to TRα (16-18). The comparison of TR mutant mice showed that the hypertrophic effect of TH is predominantly mediated by TRα action independent from DNA-binding.

Already at baseline, TRα^KO^ mice had smaller posterior and interventricular wall diameters (LVPW;d and IVS;d) and lower absolute and relative heart weights compared to their WT littermates, whereas hearts of TRβ^KO^ mice did not differ from WT mice. T3 treatment induced ventricular growth with increased wall diameters and heart weights in WT and TRβ^KO^ mice, but not in TRα^KO^ mice. Absence of TRβ had no effect on T3-induced cardiac hypertrophy, whereas absence of TRα completely abolished the T3 effect. Thus, the hypertrophic effect of T3 was mediated by TRα.

In contrast to TRα^KO^ mice, TRα^GS^ hearts were not smaller at baseline. Wall diameters, heart weight and cardiomyocyte size of TRα^GS^ mice increased significantly with T3 treatment. These results demonstrate that DNA binding is not required for TRα to mediate the hypertrophic effect of TH. One of the best characterized noncanonical effects of TRα is PI3K activation (10, 11, 30). PI3K signaling is crucial in postnatal cardiac growth and constitutively active PI3K leads to cardiac hypertrophy with increased cardiomyocyte size (31), similar to T3-treated WT and TRα^GS^ mice, further supporting the concept that noncanonical DNA-binding independent TRα action underlies T3-induced cardiac growth.

We did not observe a change in cardiac function after seven weeks of T3 treatment. This is not necessarily surprising, because even in mice with constitutively active PI3K fractional shortening was not changed (31). We studied T3 effects only on healthy hearts and hypothesize that effects of noncanonical T3 signaling may become apparent in heart failure.

A more general question is whether TH induced cardiac hypertrophy is a response to systemic activation of metabolism with increased demand of blood flow, a consequence of central activation of the sympathetic nervous system by TH or a direct, heart intrinsic effect of TH. To determine precisely where T3 and TRα act would require studies in cell-type specific TR KO and KI mice, while we here studied global TR KO and KI mice. Our results therefore do not allow to infer the site or cell-type of TH action. As our data from TR mouse models demonstrate that noncanonical TRα signaling is mainly responsible for cardiac hypertrophy and PI3K activation in mouse hearts was necessary for cardiac hypertrophy (16-18), we hypothesize that cardiac hypertrophy is a heart intrinsic effect of TH.

### Absence of TRα can be compensated by TRβ

*Myh7* encodes for a myosin heavy chain beta (MHC-β) isoform (slow twitch) expressed primarily in the heart and is repressed by TH. In contrast to untreated WT mice, basal *Myh7* expression was not repressed in TRα^KO^ mice and TRα^GS^ mice, mice without TRα-DNA-binding, which demonstrates that TRα and its canonical mode of action mediate *Mhy7* repression. T3-treatment repressed *Mhy7* expression in WT mice. Strikingly, *Mhy7* was as completely repressed in T3- treated TRα^KO^ and TRα^GS^ mice as in WT mice. This response to T3 must be independent from TRα. Our interpretation is that in the absence of TRα, TRβ substitutes for TRα and leads to *Myh7* repression. The pattern of *Hcn2* expression supports this hypothesis further: *Hcn2* was induced by T3 not only in WT mice, but also in TRα^KO^ and TRα^GS^ mice, which must again be independent from TRα, suggestive of substitution by TRβ. In TRβ^KO^ mice, T3 induced *Hcn2* and repressed *Mhy7*, which suggests that to a certain degree both TR isoforms can substitute for each other’s canonical action. This may be the case for many genes and several organs because an *in vitro* study in neural C17.2 cells found that a substantial fraction, but not all, of the T3 target genes display a preference for one of the two receptor isoforms (32). More than one third of TR binding sites were shared by both TR isoforms, suggesting that both receptors contribute to regulation of the same genes as we found here *in vivo*. *Prkcb* showed an interesting expression pattern with T3-induced induction in WT and TRα^GS^ but not TRα^KO^ mice. Its expression seems to be independent from of TRα DNA-binding, a pattern we had previously observed for TRβ^GS^- repressed genes in liver (*Scd1*, *Fasn*) (7), suggesting that noncanonical of TRα action regulates gene expression although the mechanism remains to be determined.

### Heart rate

We had previously seen in an *ex vivo* model that heart rates of hyperthyroid TRα^KO^ and TRα^GS^ mouse hearts were comparable to WT mouse hearts (33). Our interpretation was that residual TRβ expression may partially compensate for the absence of TRα. Here, *in vivo* heart rate measured with radio telemetry was again responsive to TH in global TRα^KO^ and TRα^GS^ mice. In both genotypes heart rate was shifted in hypothyroidism and with T3 treatment, although with a narrower range than in WT mice. This regulation is most likely mediated through TRβ, apparently exerting the same function as the TRα. While basal heart rate in ECG was clearly dependent on TRα, long-term comparison of hypothyroidism and T3 treatment with telemetry revealed a TH-mediated, but TRα-independent mechanism. Possibly, canonical TRα action controls basal and circadian heart rate while long-term adaption of heart rate in thyroid dysfunctions is likely to be controlled by TRβ.

## Conclusion

As gene expression (*Myh7*, *Hcn2*) and heart rate, both canonically regulated, were responsive to T3 treatment in TRα^KO^ mice we conclude that absence of canonical TRα signaling was compensated by TRβ. In contrast, T3 treatment for seven weeks could not induce cardiac hypertrophy in TRα^KO^ mice and TRβ could not substitute for this noncanonical TRα effect. We conclude that TRβ can substitute for canonical, but not noncanonical TRα signaling. This difference can be explained mechanistically. While TRα and TRβ share a large part of DNA binding sites, explaining overlapping canonical signaling, their noncanonical signaling has evolved differently after emergence of the two TR isoforms (34). For example, the SH2 binding motif in TRβ that is crucial for TRβ/p85 association and PI3K activation is not present in TRα. Consequently, noncanonical signaling seen for TRβ was absent with TRα in CHO and rat pituitary cells (6). Hence, TRα and TRβ are unlikely to be able to substitute for each other’s noncanonical signaling and this concept is supported here with *in vivo* data.

The present phenotypic observations from TR mutant mice may have implications beyond cardiac gene expression and heart rate. Currently, differences in physiological effects of TRα and TRβ are attributed to TR isoform specific target genes or the relative expression of TR isoforms in cells. Based on our current results, we suggest that the difference in noncanonical TR action between TRα and TRβ also contributes to phenotypic TRα and TRβ differences.

In conclusion, the comparison of physiological T3 effects in WT and TR mutant mice demonstrates that ventricular growth is largely mediated by noncanonical TRα action, whereas cardiac gene induction and heart rate regulation are mainly regulated by canonical TRα action. TRβ can partially substitute for absent canonical but not noncanonical TRα action.

## Author contributions

L.C.M. conceived the project. Experiments were designed by D.G., G.S.H., N.S., D.F., K.L. and L.C.M. D.G., G.S.H., S.C.G., J.P. and K.L. performed the experiments and all authors analyzed and interpreted data. H.F., V.G.-D., M.H.A. designed and supervised the heart phenotyping at the German Mouse Clinic. D.G., G.S.H. and L.C.M. wrote the manuscript and all authors contributed to the final version.

## Acknowledgements

We thank Konstanze Schättel, Kathrin Strumann, Jonas Bodmann and the staff of the IMCES facility for expert technical assistance. We are grateful for the continued dedicated support from Prof. Dr. G. Hilken, Dr. A. Wißmann, Dr. P. Dammann and the staff of the core animal facility at the University Hospital Essen.

