## Supplement for "Canonical and noncanonical contribution of thyroid hormone receptor isoforms alpha and beta to cardiac hypertrophy and heart rate in male mice"

**Supplemental information**

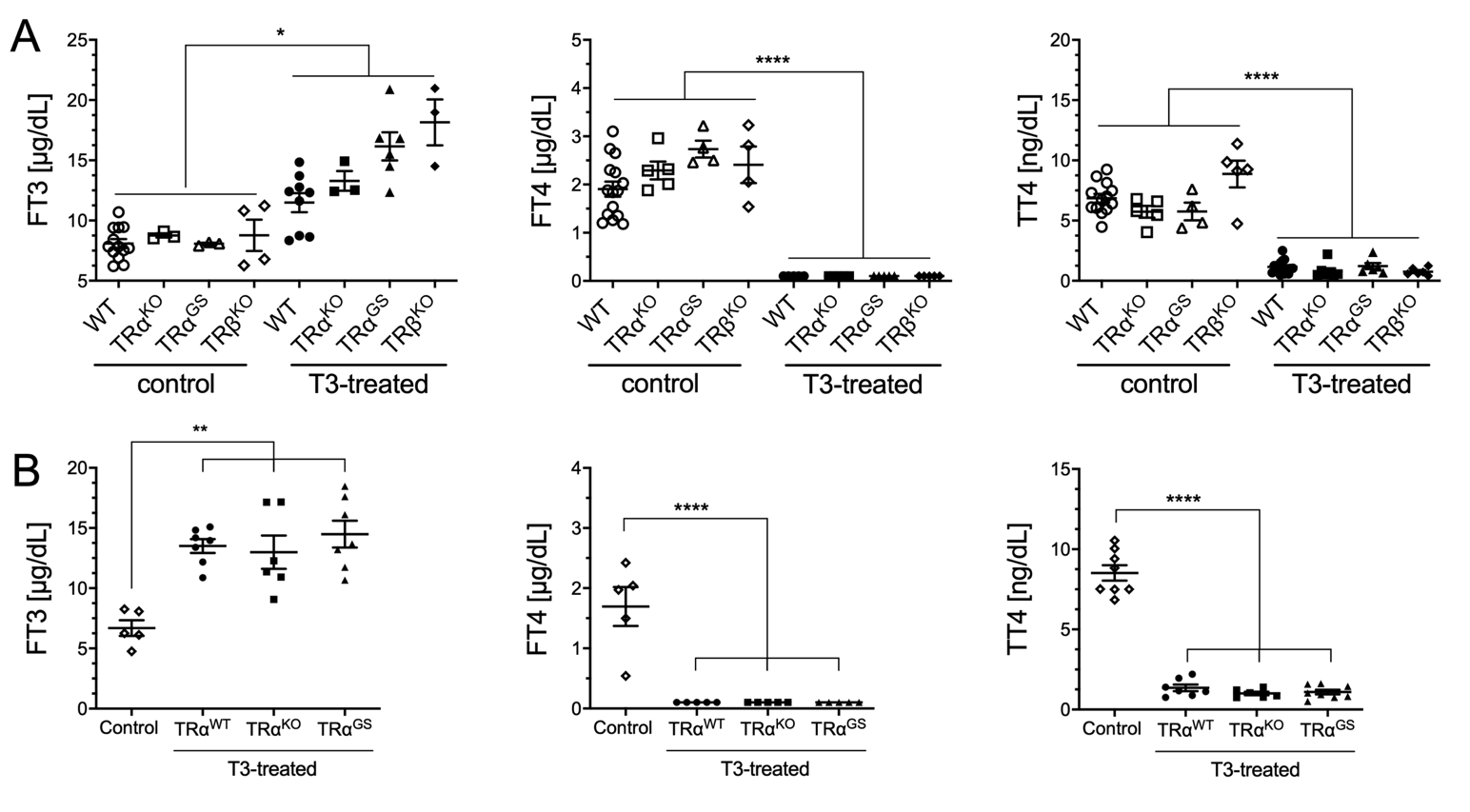

Figure S1: Thyroid hormone serum concentration of control and T3-treated mice. (A) TH serum concentrations of free T3 (FT3), free T4 (FT4) and total T4 (TT4) from control and T3-treated male WT (circles, n=9-15), TRα^KO^ (squares, n=3-5), TRβ^KO^ (triangle, =3-5) and TRα^GS^ (n=3-6) mice (mean ± SEM; *P< 0.05; one-way ANOVA with Sidak‘s multiple comparison test; control vs. T3-treated). (B) FT3, FT4 and TT4 concentrations in untreated control (diamonds, n=5-8), T3-treated TRα^WT^ (circles, n=5-7), TRα^KO^ (squares, n=5-6) and TRα^GS^ (triangles, n=5-8) mice. (Mean ± SEM; One-way ANOVA with Dunnett‘s multiple comparison test, control vs. T3-treated respectively; *P<0.05, **P<0.01, ****P<0.0001.)

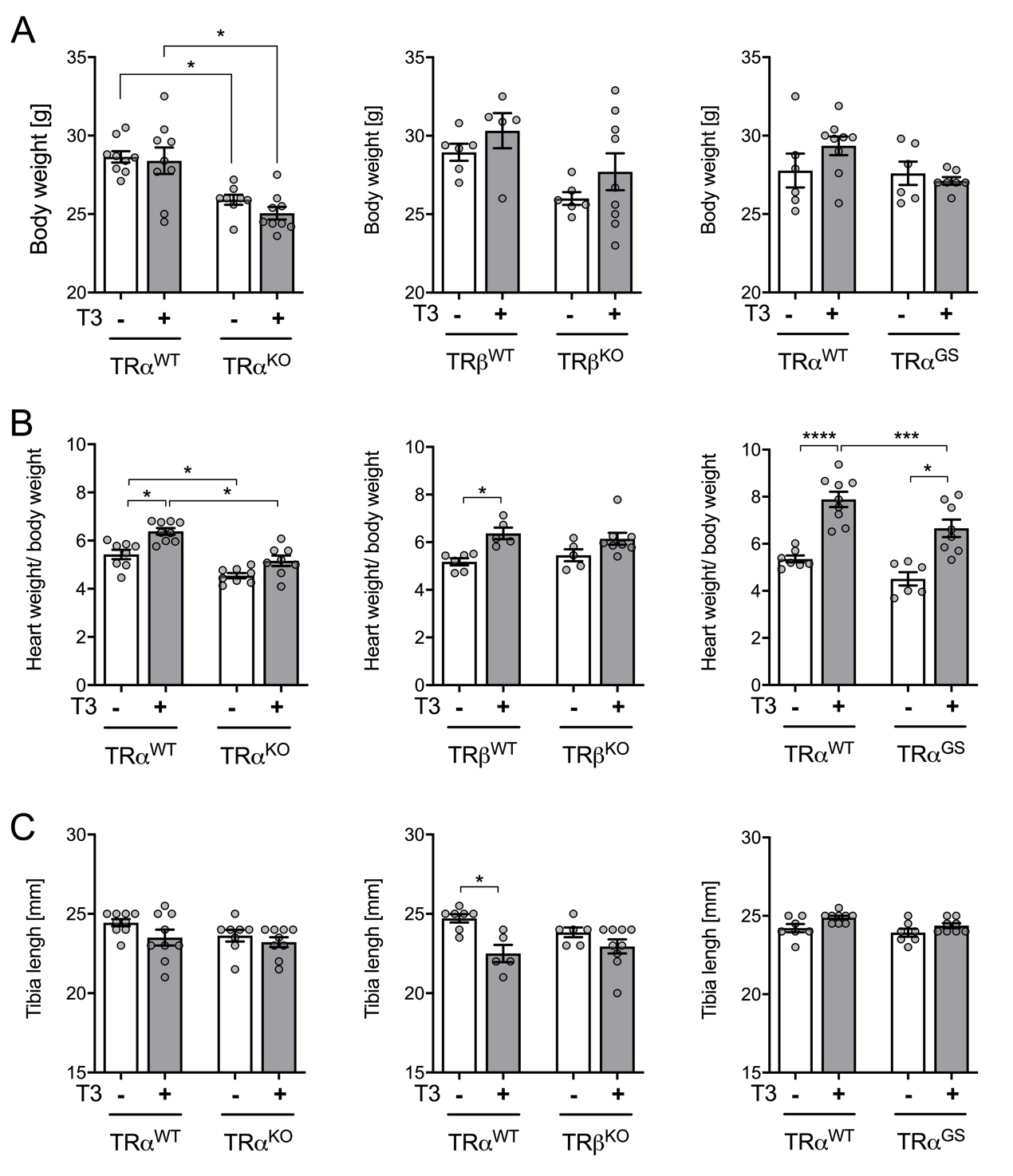

Figure S2: *Ex vivo* heart weight analysis of control and T3-treated TRα^WT^, TRα^KO^, TRβ^WT^, TRβ^KO^, TRα^WT^ and TRα^GS^ mice. (A) Body weight. (B) Heart weight normalized to body weight. (C) Tibia length. (n=5-8; mean ± SEM; one-way ANOVA with Sidak‘s multiple comparison test; *P<0.05, ***P<0.001; ****P<0.0001.)

Table S1: Primers sequences for gene expression analysis

| Gene | Forward | Reverse | Accession No. |
| --- | --- | --- | --- |
| *Polr2a* | CTTTGAGGAAACGGTGGATGTC | TCCCTTCATCGGGTCACTCT | NM_001291068 |
| *Myh6* | CAGACAGAGATTTCTCCAACCCA | GCCTCTAGGCGTTCCTTCTC | NM_010856.4 |
| *Myh7* | CACGTTTGAGAATCCAAGGCTC | CTCCTTCTCAGACTTCCGCA | NM_080728.2 |
| *Hcn2* | CCAGTCCCTGGATTCGTCAC | TCACAATCTCCTCACGCAGT | NM_008226.2 |
| *Adrb1* | GCCCTTTCGCTACCAGAGTT | ACTTGGGGTCGTTGTAGCAG | NM_007419.2 |
| *Adrb2* | ACTTCTGGTGCGAGTTCTGG | GCTCTGGTACTTGAAGGGCG | NM_007420.2 |
| *ATP2a2* | AACTACCTGGAACAACCCGC | TCATGCAGAGGGCTGGTAGA | NM_001110140. |
| *Pln* | TTCATGCTCTGCACTGTGACG | GCCAAATGTGAGCTGTCTTCTTTT | NM_001141927.1 |
| *Actc1* | TAG CAC GCC TAC AGA ACC CA | GGATACCTCGCTTGCTCTGG | NM_009608.4 |
| *Bcl3* | CTGAACCTGCCTACTCACCC | AGTATTCGGTAGACAGCGGC | NM_033601.3 |
| *Bax* | TTTTGCTACAGGGTTTCATCCAG | TTCATCTCCAATTCGCCGGA | NM_007527.3 |
| *Prkcb* | GGATTCCAGTGTCAAGTCTGCT | GGTCACAGAAGG TAG CA | NM_001316672.1 |
| *Pkb* | GGATTCCAGTGTCAAGTCTGC T | GGTCACAGAAGGTAGCA | NM_008855.2 |
| *18s* | CGGCTACCACATCCAAGGAA | GCTGGAATTACCGCGGCT | NR_003278.3 |
| *Ppia* | CTTGGGCCGCGTCTCCTTCG | GCG TGTAAAGTCACCACCCTGGC | NM_008907.7 |
| *Gapdh* | CCTCGTCCCGTAGACAAAATG | TGAAGGGGTCGTTGATGG C | NM_001289726.1 |
